# Human Cytomegalovirus Remodeling of Lipid Metabolism Requires pUL37×1

**DOI:** 10.1101/526228

**Authors:** John G. Purdy, Yuecheng Xi, Samuel Harwood, Lisa Wise

**Affiliations:** University of Arizona, Department of Immunobiology, Tucson, Arizona, USA; University of Arizona, BIO5 Institute, Tucson, Arizona, USA; University of Arizona, Department of Molecular and Cellular Biology, Tucson, Arizona, USA

## Abstract

Human cytomegalovirus (HCMV) replication requires remodeling of the host metabolic network, including increasing central carbon metabolism, fatty acid elongation and lipid synthesis. The virus-host interactions regulating HCMV metabolic hijacking are essential to infection. While multiple host factors including kinases and transcription factors have been defined, little is known about the viral factors required. Some host factors involved are dependent on Ca^2+^ signaling or host stress response. The viral pUL37×1 protein mobilizes Ca^2+^ and targets the mitochondria. The Ca^2+^ changes induced by pUL37×1 can activate stress responses. In this study, we tested the hypothesis that the viral pUL37×1 protein is required for HCMV metabolic remodeling. Using a combination of metabolomics, lipidomics, and metabolic tracers, we demonstrate that pUL37×1 protein is necessary for HCMV induced fatty acid elongation and lipid metabolism but not required for most viral hijacking of central carbon metabolism. Additionally, we show that pUL37×1 is required for HCMV to fully induce two host proteins that were previously demonstrated to be important for virus induced lipid metabolism: fatty acid elongase 7 and the ER-stress related kinase PERK. We also demonstrate, for the first time, that HCMV replication results in the increase in phospholipids with very-long chain fatty acyl tails. We conclude that pUL37×1 is required for HCMV metabolic remodeling but is not necessary to for the virus to hijack metabolism on a network-wide scale.

**Importance:** Human cytomegalovirus (HCMV) is a common pathogen that infects most people and establishes a lifelong infection. Infection is generally asymptomatic. However, HCMV can cause end-organ disease that results in death in the immunosuppressed. Additionally, HCMV is the leading cause of congenital infection and birth defects. Children born with congenital HCMV may suffer lifelong disabilities such as hearing loss, microcephaly, and developmental impairment. Although there are treatments approved for HCMV, none offer a cure. HCMV infection depends on remodeling host metabolism. Blocking the virus from hijacking metabolism limits HCMV replication. However, the viral mechanisms for remodeling metabolism are poorly understood. In this study, we demonstrate that the viral pUL37×1 protein is required for HCMV to increase fatty acid elongation and remodel lipid metabolism. Our findings define an important viral player in controlling the virus-host metabolism interaction.

## Introduction

Human cytomegalovirus (HCMV) is a β-herpesvirus that established a persistent lifelong infection. Infection is common, a majority of people are infected (1). HCMV is an opportunistic pathogen that can cause severe and life-threatening diseases when the immune system is compromised, such as in solid-organ and stem cell transplant patients or those with HIV/AIDS (1,2). HCMV is also a major cause of birth defects (2). Congenital infection can result in hearing loss, microcephaly, developmental disabilities, and fetal loss (3,4). HCMV infections have been associated with cardiovascular disease (5,6), cancers (7) and age-related immune function (8-10).

Since HCMV—like all viruses—does not encode a metabolic network, it relies on the host to provide the energy, metabolites, and lipids required for replication. Limiting nutrients or targeting metabolic pathways inhibits HCMV replication (11-16). Virus replication results in a global change in metabolism, altering the concentration of most metabolites that have been examined (11,12,14-24). HCMV infection alters central carbon metabolism and increases the utilization of glucose and glutamine (11, 20,23, 25-27). Infection increases the flow of carbons from glucose to lipid synthesis (11, 12,17,24,28-30). HCMV incorporates specific lipids into its envelope (12,31). Importantly, lipids synthesized during infection are required to build the virus envelope membrane (12). Previously we demonstrated that carbons from glucose are used for fatty acid (FA) elongation to generate very long-chain fatty acids (VLCFA) through the action of fatty acid elongase 7 (ELOVL7) (12,17). ELOVL7 is required for efficient virus release and virion infectivity, demonstrating the fatty acid elongation is required for infection (12).

The virus-host metabolism interactions involve various host and viral factors. HCMV replication depends on AMPK dependent metabolic regulation (21,32). During infection, HCMV actives AMPK through Ca^2+^-calmodulin-dependent kinase kinase (CaMKK) activity (32). CaMKK is required for HCMV hijacking of glycolysis (22). HCMV limits AMPK downregulation of FA synthesis (33). HCMV induced ELOVL7 activity requires the early viral protein pUL38 (12). pUL38 prevents mammalian target of rapamycin (mTOR) inactivation and induces the maturation of sterol regulatory element binding proteins (SREBPs) transcription factors to enhance FA metabolism (12,33,34). In addition to mTOR, other kinases are also used by HCMV to hijack metabolism. The endoplasmic reticulum (ER) stress kinase, PKR-like ER kinase (PERK, also known as eukaryotic translation initiation factor 2-alpha kinase 3, EIF2AK3) is also activated by HCMV to increase lipid synthesis (29).

Although some of the host factors that HCMV uses to modulate specific metabolic pathways have been defined, little is known about the viral factors that required to coordinate the hijacking of the host metabolic network beyond our initial understanding of the role of pUL38. Since CaMKK activity is stimulated by Ca^2+^, viral stimulation of glycolysis may depend on intracellular Ca^2+^ mobilization. HCMV infection increases intracellular Ca^2+^ concentrations (35). HCMV pUL37×1 protein triggers the release of ER stored Ca^2+^ into the cytosol (36). Thus, pUL37×1 mediated increase of Ca^2+^ into the cytosol could alter metabolism (Fig. 1). In addition to Ca^2+^ release, pUL37×1 blocks apoptosis and targets the mitochondria (37-40) while increasing the fragmentation of the mitochondrial network (41). Given the importance of the mitochondria to metabolism, this could be an additional mechanism for HCMV metabolic control (Fig. 1). pUL37×1 traffics to the outer mitochondrial membrane where it contacts the ER (42). The subcellular location of pUL37×1 or stress-associated with the release of stored ER Ca^2+^ may influence ER-related metabolism, such as FA elongation and lipid synthesis.

**Figure 1.**
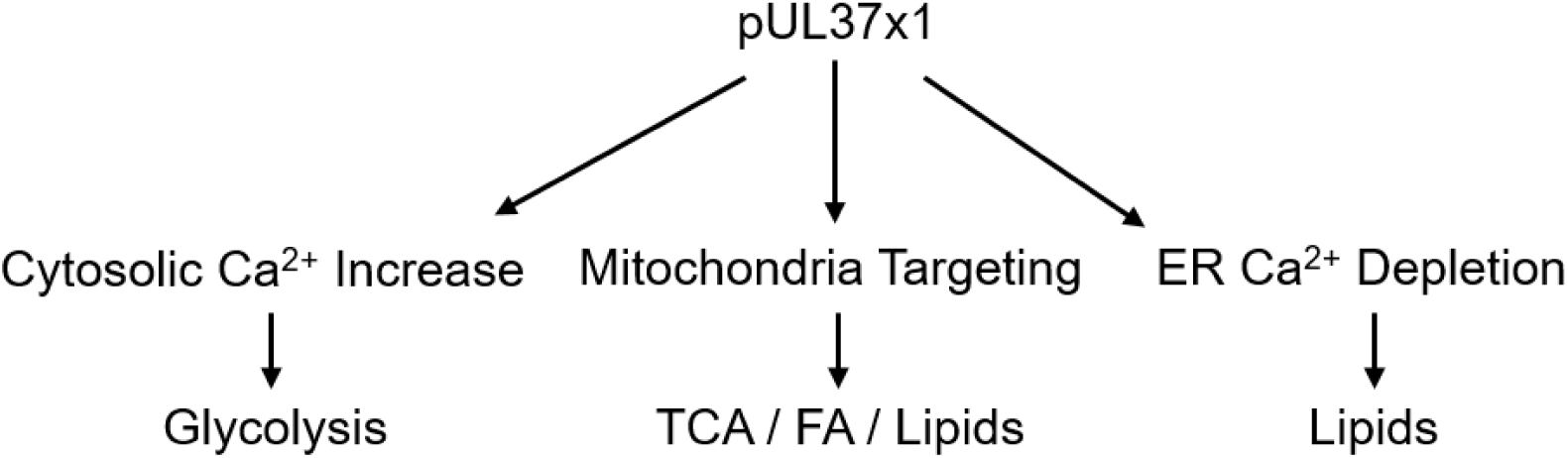
Model for possible metabolic regulation imposed through the various functions of pUL37×1. HCMV induced increase in cytosolic Ca^2+^ could induce a Ca^2+^ dependent signaling cascade that leads to an increase in glycolytic activity. pUL37×1 localization to the mitochondria may affect mitochondrial metabolic pathways including TCA, FA and lipid metabolism. A third possibility is the depletion of Ca^2+^ from the ER by pUL37×1 may alter ER localized metabolism such as lipid synthesis or ER-stress related metabolic activity. Greater details are described in the text.

Given pUL37×1 function in Ca^2+^ mobilization and localization to membranes important for metabolism, we—and others (18,22,43,44)—hypothesize that HCMV metabolic remodeling requires pUL37×1. We tested this hypothesis using a mutant virus that lacks the UL37×1 gene (36,45) in combination with metabolomics, lipidomics and metabolic tracing experiments. We found that pUL37×1 is required for the overall robustness of HCMV metabolic remodeling; however, the virus is able to induce metabolic hijacking independent of pUL37×1. Specifically, pUL37×1 supports HCMV induction of FA and lipid metabolism while increasing ELOVL7 and PERK protein levels. Moreover, our findings establish that HCMV infection results in a robust increase in phospholipids with very long chain fatty acid (PL-VLCFA) tails. pUL37×1 is important for the dramatic increase in these PL-VCFAs. Additionally, pUL37×1 suppresses the levels of phosphatidylserine lipids with shorter tails that are associated with apoptotic signaling. The findings reported here improve our understanding of the viral mechanisms used by HCMV to remodel metabolism and further illustrates that HCMV commandeers metabolism to generate a metabolic environment and lipidome that supports infection.

## Results

Active HCMV replication requires the products of various metabolic pathways including those involved in energy, amino acid, nucleotides, fatty acids and lipids. HCMV pUL38 prevents mTOR deactivation and stimulates SREBP maturation and fatty acid elongation (12,33). Beyond pUL38, we have a limited understanding of the viral mechanisms used to alter varied metabolic pathways. We and others (43) hypothesize that pUL37×1 is required for HCMV metabolic control. We directly tested this hypothesis by comparing the viral control of metabolite concentrations in WT infected cells to those infected with a mutant virus that lacks the UL37×1 gene (36).

### Global metabolome remodeling by HCMV occurs largely independent of pUL37×1

Our work builds on the well-defined HCMV remodeling of the metabolome of primary human fibroblast cells under fully-confluent, serum-free conditions (11, 12, 14-17,22, 23, 32, 33). Since most prior characterization of pUL37×1-null viruses was done under sub- or near-full confluent conditions in the presence of animal serum, we briefly characterized the mutant virus infection in confluent, serum-free conditions. We confirmed that our pUL37×1-null mutant virus, *sub*UL37×1 (36), expresses pUL36 and pUL38 while ablating pUL37×1 expression (Fig. 2A). The cytopathic effect (CPE) characterized by WT infection—e.g. visual cytomegaly—was lessened in cells infected with the *sub*UL37×1 mutant (Fig. 2B). At the later stages of infection, a greater amount of cellular debris was observed in mutant virus infected cells, consistent with an increase in cell death at 3-4 days post infection (dpi) (Fig. 2B). We observed a strong loss in virus replication with UL37×1 deletion (Fig. 2C). Furthermore, the loss of pUL37×1 also resulted in the release of fewer infectious virions per total released viral particles (Fig. 2D). Overall, these results were consistent with observations of UL37x-1 null mutant viruses in sub-confluent cells grown in serum (36,38,39).

**Figure 2.**
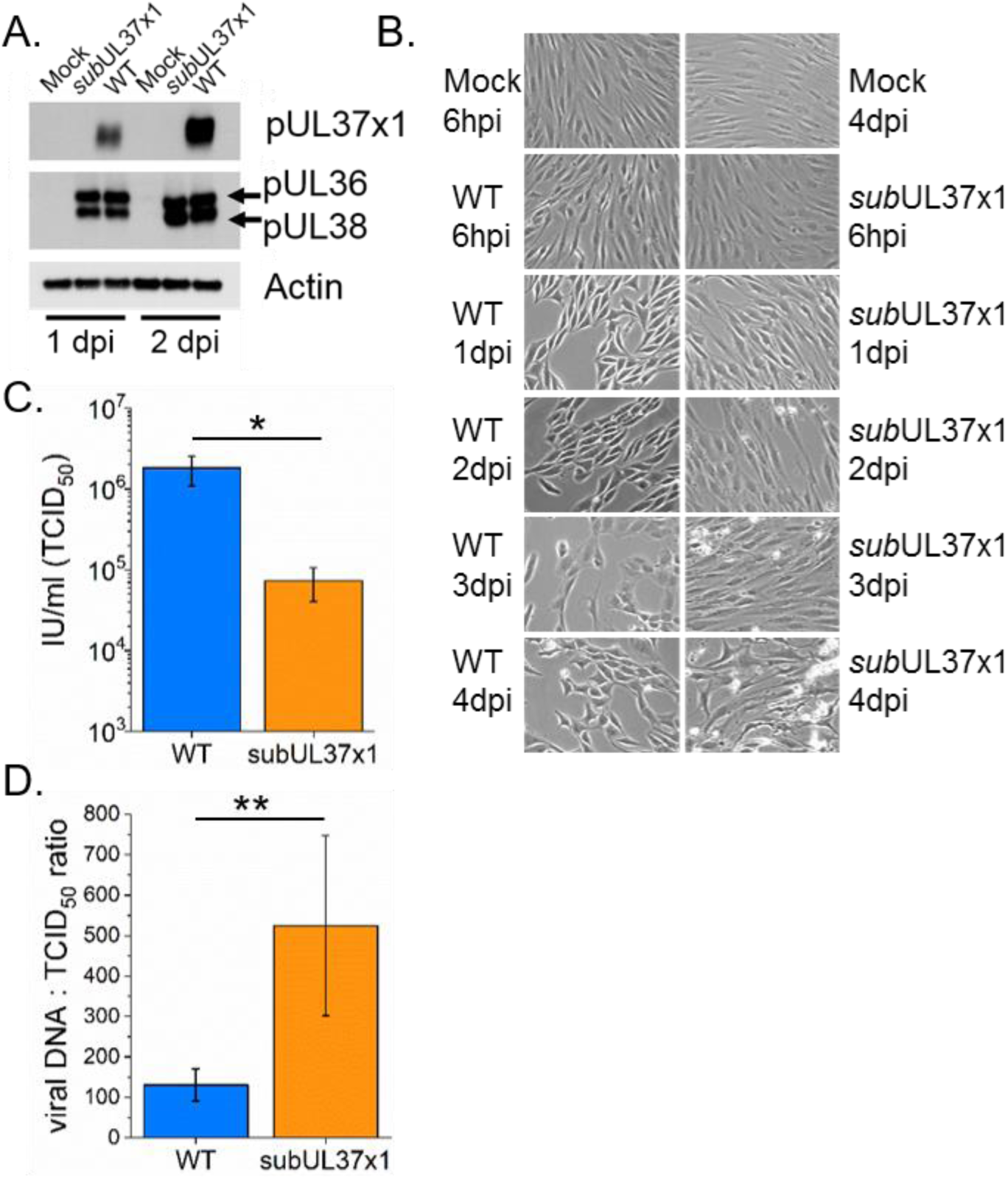
UL37×1 is required for cell survival and virus replication in serum-free conditions. (A) Western blot analysis reveals that *sub*UL37×1 fails to express pUL37×1 while the expression of neighboring genes is unaffected. (B) Infection in fully confluent fibroblast cells in serum-free conditions was visually tracked from 6 hpi to 4 dpi. Images from uninfected control cells at 6 hpi and 4 dpi is also included. (C) Infectious virus particles released by cells infected with WT or mutant *sub*UL37×1 was measured at 4 dpi at MOI = 3. (D) The particle-to-infectious units as measured by viral DNA and TCID50 were compared for particles released by WT or subUL37×1 infected cells at 4 dpi, MOI = 3. For (C) and (D) the error bars are the standard deviation from three independent experiments (N = 3). * p = 0.05 – 0.01; ** p = <0.01; t-test.

Next, we tested if pUL37×1 is required for viral hijacking of the host metabolome using liquid-chromatography high-resolution tandem mass spectrometry (LC-MS/MS). Infection with WT AD169 virus resulted in a global remodeling of metabolites that were most dramatically changed at 1 dpi and later (Figure 3A). HCMV strongly increases glycolytic and TCA cycle metabolites along with others that have been previously described (11, 14, 15, 19, 21, 22, 46). Infection with the mutant *sub*UL37×1 virus similarly altered metabolite levels starting at 1 dpi (Figure 3A). The correlation between metabolic profiles of WT and *sub*UL37×1 infected cells was low early in infection (Fig. 3B, 6 hpi) but strongly correlated at later time points (Fig. 3B, 1-3 dpi).

**Figure 3.**
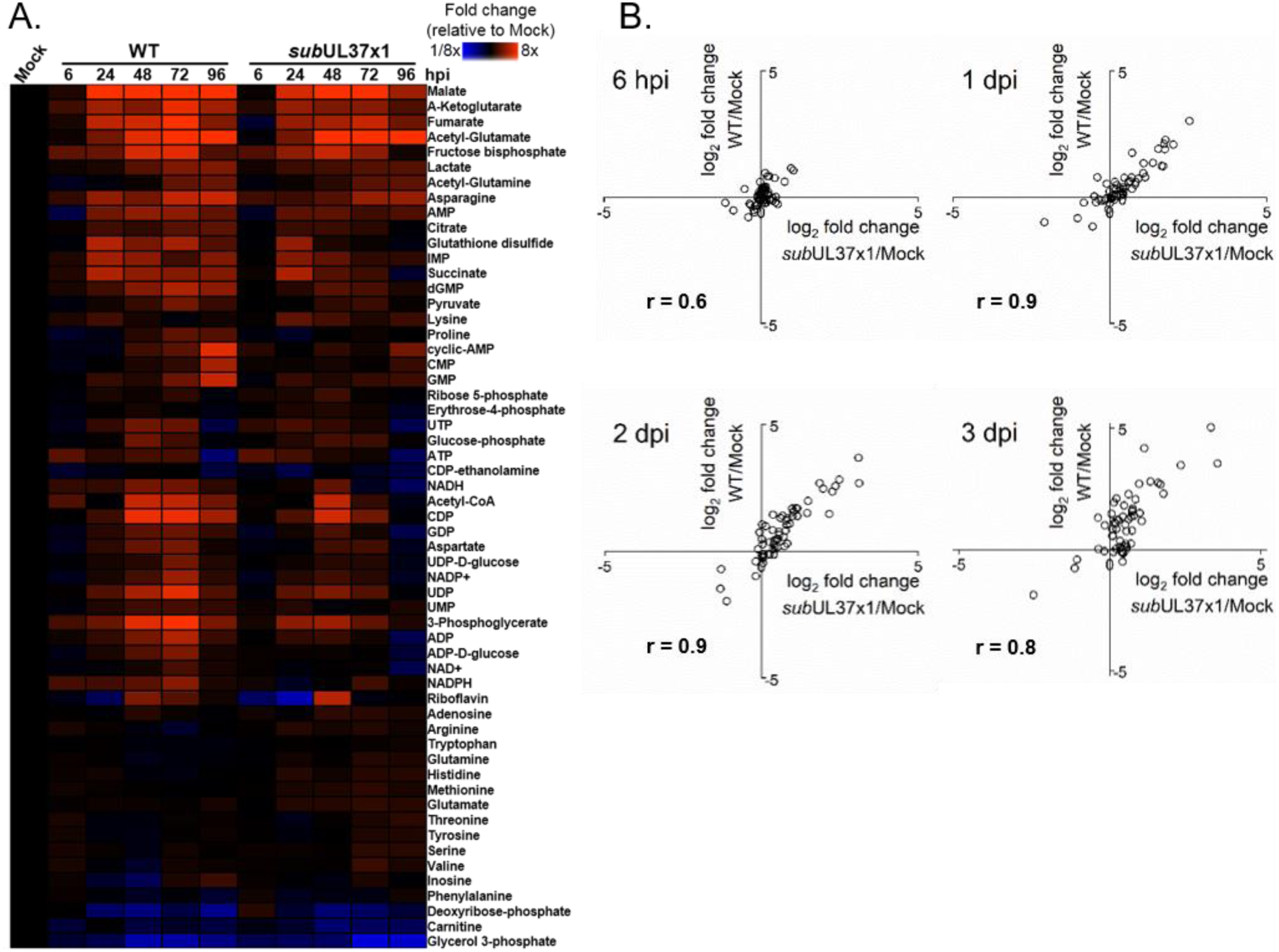
Global metabolic hijacking by HCMV occurs independent of UL37×1. (A) Metabolite levels during the course of productive replication in fibroblast cells comparing WT to mutant subUL37×1 infected fibroblast cells, MOI = 3. Fold-change in metabolite concentrations relative to uninfected cells at each timepoint (6, 24, 48, 72, and 96 hpi). (B) Correlation plots for metabolite levels normalized to uninfected cells (WT = y-axis and subUL37×1 = x-axis). Correlation R values are given for each timepoint from 6-72 hpi. For both A and B, metabolites were measured from technical replicates from 4 independent experiments (N = 4).

Importantly, glycolytic and TCA metabolites—e.g., hexose-phosphate, pyruvate, malate, α-ketoglutarate, and succinate—were increased in *sub*UL37×1-infected cells compared to uninfected cells but were slightly reduced compared to WT-infected cells (Fig. 3A) These findings demonstrated that pUL37×1 may contribute to HCVM remodeling of specific metabolic pathways, but is not required for the global hijacking of the host metabolism.

### pUL37×1 is necessary for HCMV-induced FA elongation

Our metabolomic studies examined water-soluble metabolites. Hydrophobic compounds including lipids and fatty acids were not included in this type of analysis. HCMV enhances FA synthesis and dramatically increases FA elongation (11,12,17). FA synthase (FAS) forms long-chain FAs up to 16 carbons in length (C16) by connecting carbons, two at time. The product of FAS is 16-carbon saturated FA palmitate (C16:0, the number following the colon represents the number of double bonds in the FA). Palmitate can be processed further to make longer chains and/or desaturated to introduce a double bond among the carbons in the tail. In humans, longer FAs are made by one or more of the seven FA elongases (ELOVL1-7), again adding one 2-carbon unit per reaction cycle. We have previously shown that HCMV significantly increases the concentration of saturated very-long chain FAs (VLCFA), specifically those with 24 or more carbons (≥C24) (12,17).

To determine if pUL37×1 is important for HCMV induced FA metabolism, we measured FAs extracted from lipids of infected and mock-infected control cells. In cells infected with WT virus, saturated VLCFAs were increased by as much as 9-fold compared to uninfected cells confirming our previous observations (Fig. 4) (12,17). While *sub*UL37×1 virus increased some FAs compared to uninfected cells, the mutant virus failed to induce WT-like levels of VLCFAs at 2 and 3 dpi (Fig 4).

**Figure 4.**
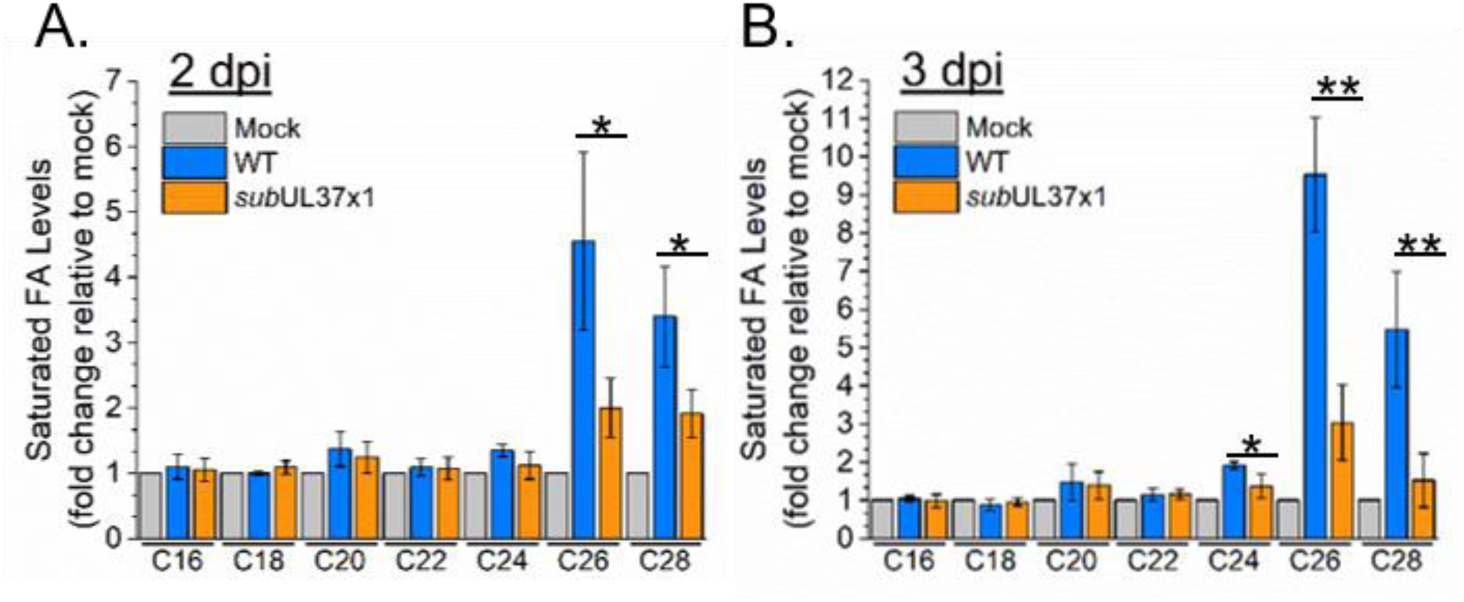
HCMV UL37×1 is required for virus induced very long chain fatty acid (VLCFA) increases. FA levels relative to uninfected cells are shown for cells infected at MOI = 3. (A) shows FA changes at 2 dpi while (B) shows changes at 3 dpi. All data represented as mean + SD of N = 4. * p = 0.05 – 0.01; ** p = <0.01; t-test.

FA elongation can be monitored by measuring the incorporation of ^13^C from labeled glucose into FAs using LC-MS, i.e., non-tandem MS1-only analysis (12,17,47). Following a 1h infection period, we fed WT and *sub*UL37×1 infected cells medium that contained ^13^C-labeled glucose at positions 1 and 6 (1,6-^13^C-glucose). Glycolytic breakdown of this labeled form of glucose will result in 1-labeled pyruvate (Fig. 5A). Pyruvate can be further metabolized generating 1-labeled citrate in the mitochondria. Citrate can be exported into the cytoplasm where it is broken down into 1-labeled acetyl-CoA and unlabeled oxaloacetate. Alternatively, pyruvate may breakdown into acetate that can be converted to acetyl-CoA (24,48). The conversation of acetyl-CoA to malonyl-CoA by acetyl-CoA carboxylase 1 (ACC-1) is the rate-limiting step of fatty acid synthesis and elongation. In our labeled cells this will result in 1-labeled malonyl-CoA per 2-carbon unit used for FA synthesis and elongation.

**Figure 5.**
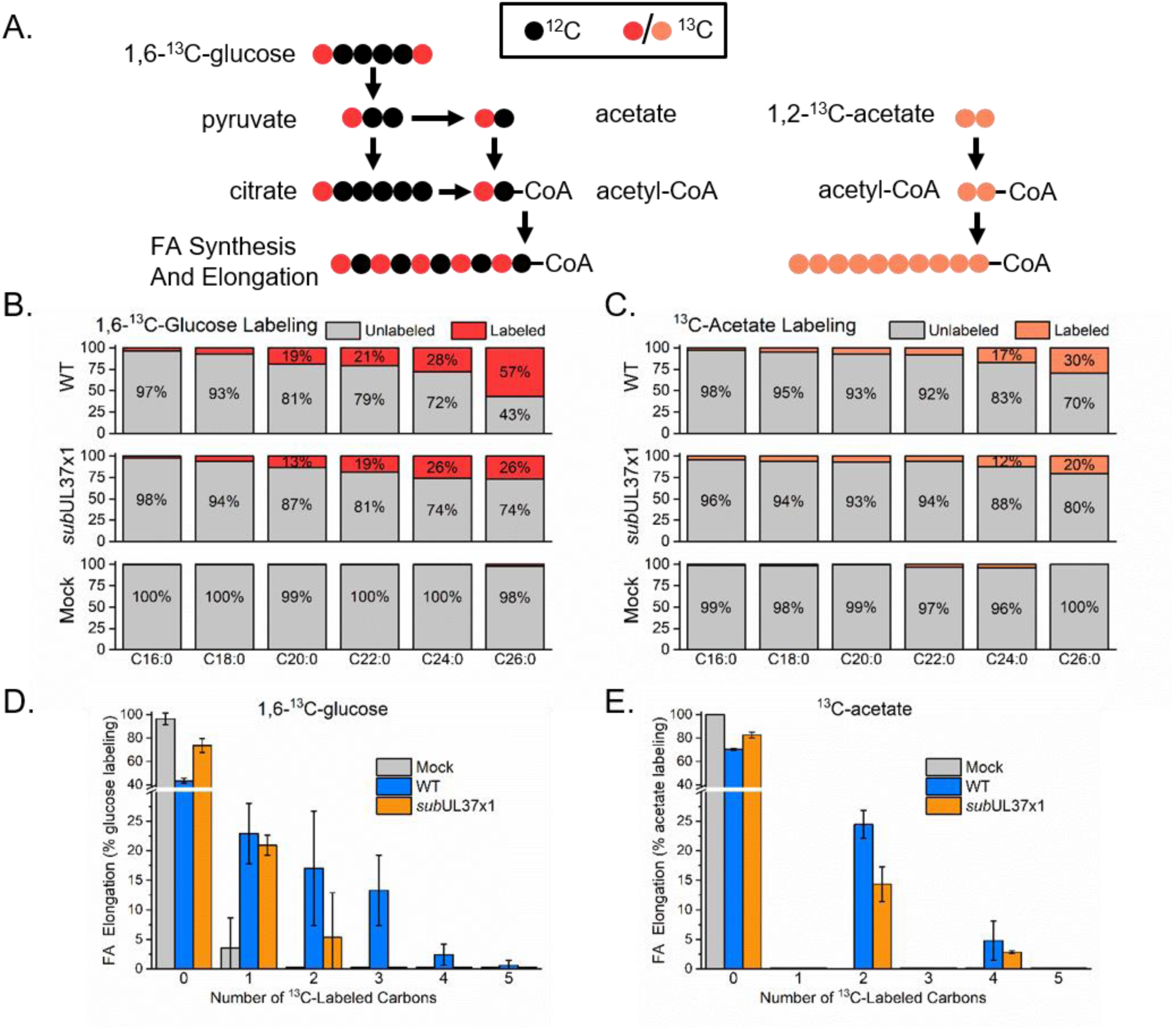
UL37×1 is needed for HCMV induced FA elongation. (A) Diagram depicting the flow of carbons from glucose and acetate to FA synthesis and elongation. Carbon atoms are represented by circles, red and orange circles represent ^13^C-labeled carbons while black circles depict unlabeled atoms. (B) Incorporation of labeled atoms from 1,6-^13^C-glucose into saturated FAs. The data is presented as the percent labeled (red) to unlabeled (gray). (C) Percent labeling of saturated FAs from 1,2-^13^C-acetate. (D-E) Labeling pattern of C26:0 from labeled glucose and labeled acetate, respectively.

Using LC-MS to measure the overall percent of FA tails that have one or more ^13^C atoms, we observed that uninfected cells have minimal to no labeling of FAs after correcting for the natural isotopic abundance of ^13^C. This is consistent with the cells being in a non-growing state due to contact-inhibition and serum-starvation. HCMV infection, however, strongly induced FA anabolism. In WT-infected cells a significantly higher proportion of FAs were labeled, and the greatest labeling occurred in longer FAs (Fig. 5B). *Sub*UL37×1 also induced FA metabolism compared to uninfected cells (Fig. 5B), however the labeling rate was less than that observed in WT-infected cells. For example, more than half of the C26:0 was labeled during WT-infection while only a quarter was labeled in *sub*UL37×1-infected cells. In addition to a reduction in the percent of labeling, the FAs isolated from mutant virus infected cells had fewer number of ^13^C atoms (Fig. 5D). In WT infected cells, we were able to observe up to 4-5 labeled atoms (or the incorporation of 8-10 new carbons). This observation was similar to that observed in cells fed uniformly labeled ^13^C-glucose where two ^13^C atoms are added per 2-carbon units (12,17). However, *sub*UL37x-1 infection resulted in only 1-2 labeled atoms indicating that only one to two 2-carbon units were added.

Our glucose labeling strategy measured the carbon contribution of both citrate and acetate that is derived from glucose. Since HCMV infection increases the contribution of acetate feeding into lipid synthesis (24), we measured the contribution of acetate to HCMV-induced FA elongation using ^13^C-labeled acetate. In this case both atoms of the 2-carbon unit were labeled. Again, cells were fed labeled acetate following a 1h infection period. Similar to the labeled-glucose tracer, only minimal labeling from acetate was observed in uninfected cells (Fig. 5C). Infection increased acetate derived atoms incorporated into VLCFAs (Fig. 5C). In WT-infected cells 30% of C26:0 was labeled from ^13^C-acetate versus 60% labeling from glucose. Additionally, only 4 labeled atoms from two 2-carbon units were incorporated (Fig 5E). The *sub*UL37×1 mutant virus induced FA elongation from acetate, but to a lower level than WT (Fig. 5C and 5E). Together our results reveal that pUL37×1 contributed to the ability of HCMV to induce FA elongation.

During infection, saturated VLCFAs are produced by ELOVL7 (12). HCMV increases the gene and protein expression of ELOVL7 (12). We examined if HCMV-induced ELOVL7 expression requires pUL37×1. In HFF cells, WT virus induced ELOVL7 expression by 2 dpi (Fig. 6), similar to our observations in MRC-5 primary fibroblasts (12). Mutant *sub*UL37×1 virus also induced ELOVL7 expression, however at a reduced level compared to WT (Fig. 6). This observation is consistent with a reduced ability of the mutant virus to induced FA elongation and the production of saturated VLCFAs.

**Figure 6.**
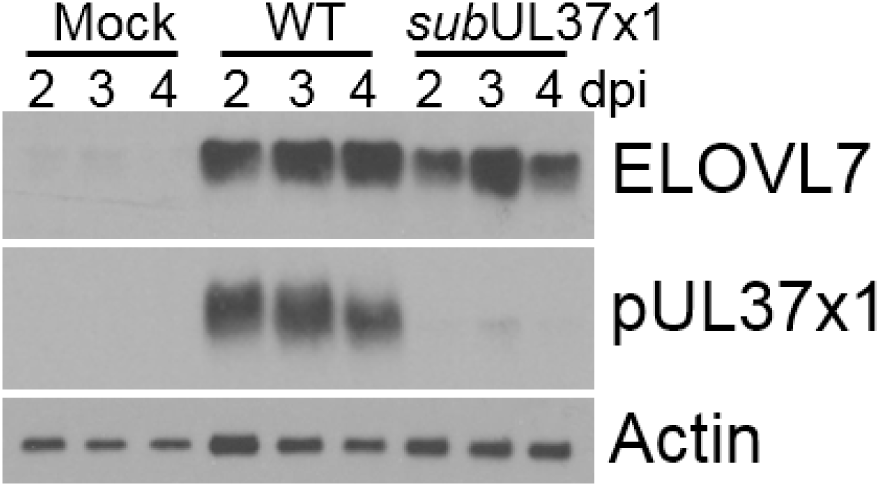
UL37×1 is necessary for full induction of ELOVL7 fatty acid elongase protein expression by HCMV. ELOVL7 protein expression was determined by Western blot analysis of uninfected, WT and mutant virus infected fibroblast cells. MOI = 3.

The pUL37×1 dependent FA synthesis of VLCFAs was consistent with a decreased ability of the *sub*UL37×1 mutant virus to induce ELOVL7 protein expression. However, pUL37×1 may also affect other aspects of FA metabolism. In addition to ELOVL7, HCMV increases the protein expression of ACC-1 and its phosphorylated form that reduces its activity (33). We found that *sub*UL37×1 also induced both forms of ACC-1 over mock levels (Fig. 7). However, there was a slight decrease in total ACC-1 levels compared to WT (Fig. 7), consistent with a decrease in FA elongation. Since the mutant virus has a strong defect in elongation from glucose derived carbons we examined if pUL37×1 influences the expression of ATP citrate lyase (ACLY) that converts citrate to acetyl-CoA in the cytosol. At late, time points in replication ACLY protein levels are slightly higher in infected cells (28). We found that ACLY levels were similar in WT and *sub*UL37×1 infected cells at 2-4 dpi (Fig. 7). Likewise, the ACSS2 enzyme that synthesizes acetyl-CoA from acetate was unaffected by pUL37×1 (Fig. 8).

**Figure 7.**
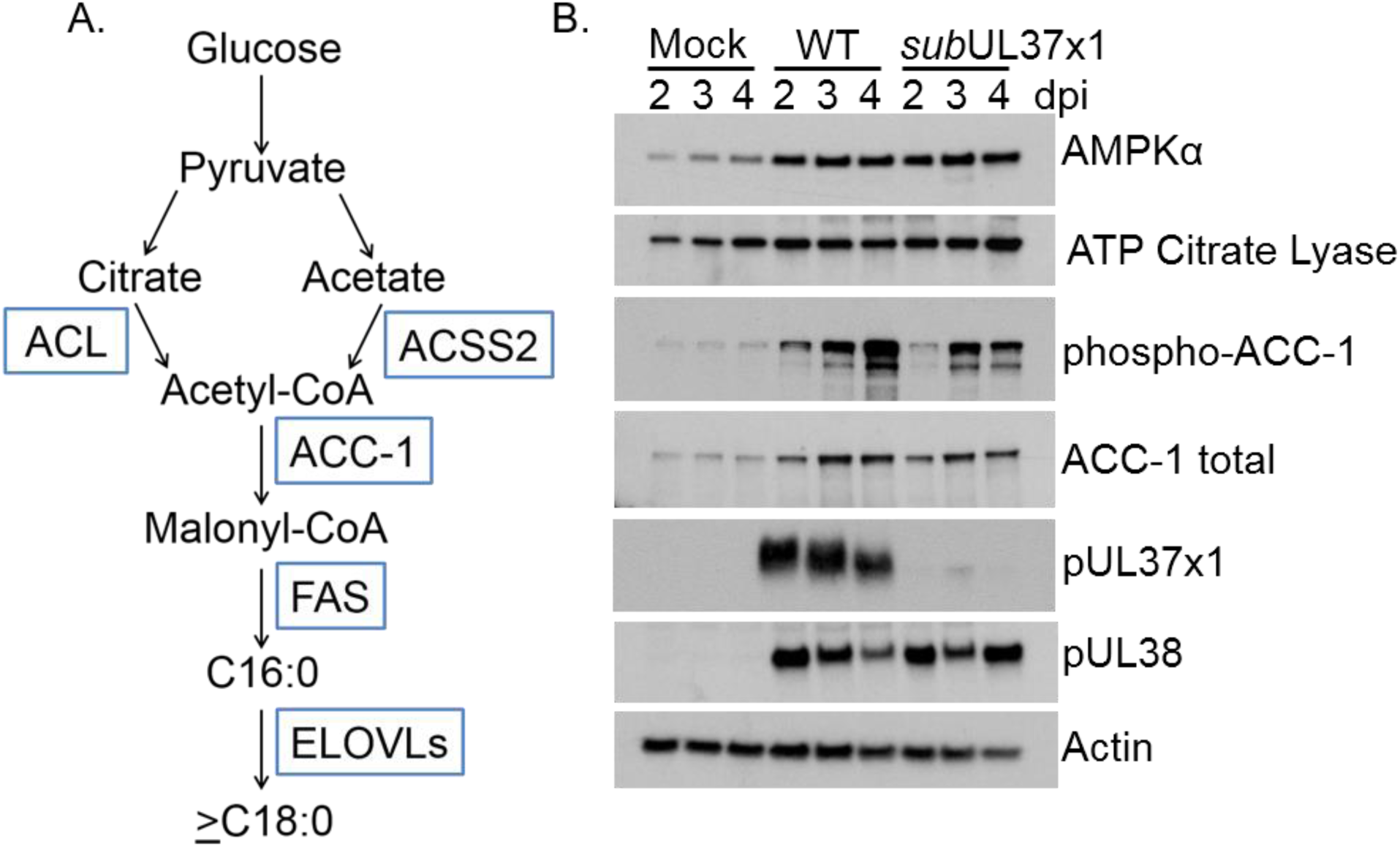
HCMV impact on proteins controlling FA synthesis. (A) Enzymes involved in metabolizing glucose carbons to FAs are shown in blue boxes. (B) Western blot of proteins involved in glucose metabolism and FA synthesis.

**Figure 8.**
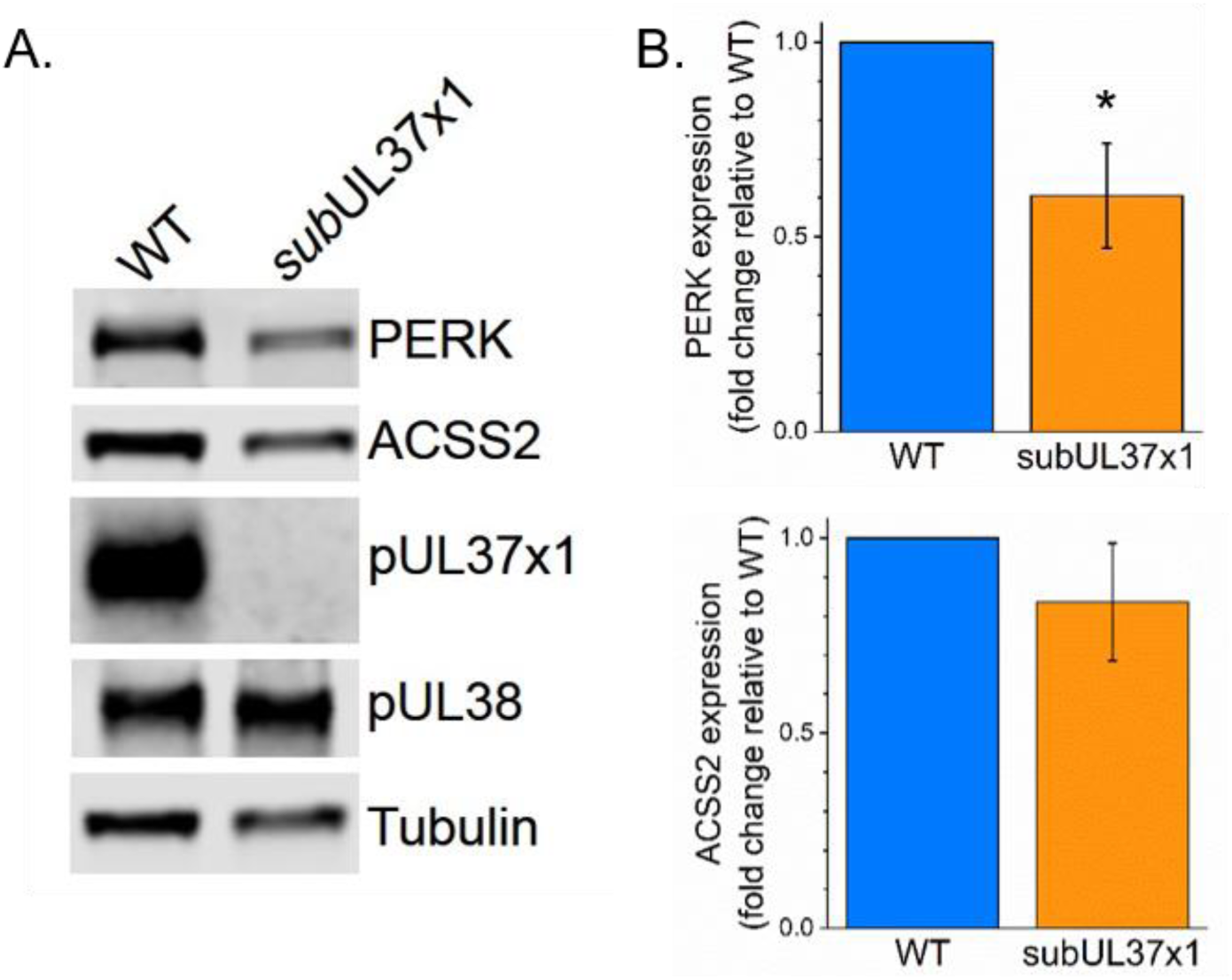
pUL37×1 is necessary for HCMV to induce PERK protein levels. (A) Western blot at 3 dpi showing PERK and ACSS2 expression level in human fibroblast cells infected with WT or mutant subUL37×1 virus at MOI = 3. (B) Quantification of PERK and ACSS2 protein levels from three independent experiments. Protein loading was normalized to the tubulin signal. * p < 0.05 using a one-sample t test.

HCMV-induced metabolic remodeling requires AMPK. HCMV induces AMPK expression and activity to increase glycolysis (21,32). However, AMPK typically phosphorylates ACC-1 to reduce its activity. We found that s*ubU*L37×1 induced the expression of the catalytic subunit of AMPK, AMPKα, to WT-like levels suggesting that pUL37×1 is not acting through AMPK mediated activity (Fig. 7). In conclusion, pUL37×1 is required for HCMV induced FA metabolism, and induction of ACC-1 and ELOVL7 protein expression.

### Viral remodeling of host lipidome requires pUL37×1

One function of FAs is to act as hydrophobic tails in lipids. Given our observations in Figs. 4-5, we hypothesized that pUL37×1 is required for the synthesis of lipids with VLCFA tails. To test this hypothesis, we developed LC-MS/MS lipidomic methods to examine phospholipids, including those that contain C24 and C26 VLCFAs. We found that WT HCMV infection dramatically increased phospholipids, especially those with VLCFA tails (PL-VLCFAs) (Fig. 9A, left panel). Phosphatidylcholines (PC) were particularly increased (Fig. 9A-B). A previous lipidomic study demonstrated that PC lipid species with long chain fatty acid tails were increased by HCMV, while those with shorter chains decreased (31). Notably, this previous analysis only examined lipids with a total of 30-42 carbons in their tail. Since the identity of any individual tail was not reported it is unlikely that this analysis included ≥C26 VLCFAs. Using our LC-MS/MS methods, we examined lipids with up to 48 total carbons in their tails. HCMV induced the levels of several PC species with 44-48 carbons in their tails (Fig. 9B). Of the top six PC species increased by HCMV, their levels were increased 17-32-fold (PC(48:7)) to 270-14,00-fold (PC46:1)). In this case, the PC tails contain a total of 48 carbons and 7 double bonds and 46 carbons and a single double bond, respectively. Since the number of carbons and double bonds impart a large amount of diversity observed in lipidomes, we examined the tail composition of these six PC species (Fig. 9D). All contained a ≥C26 tail. The greatest diversity was observed in PC(44:1). We found evidence of four different PC(44:1) species: C18:1+C26:0, C18:0+C26:1, C16:0+28:1, and C14:0+C30:1. The C30 tail was the longest one observed.

**Figure 9.**
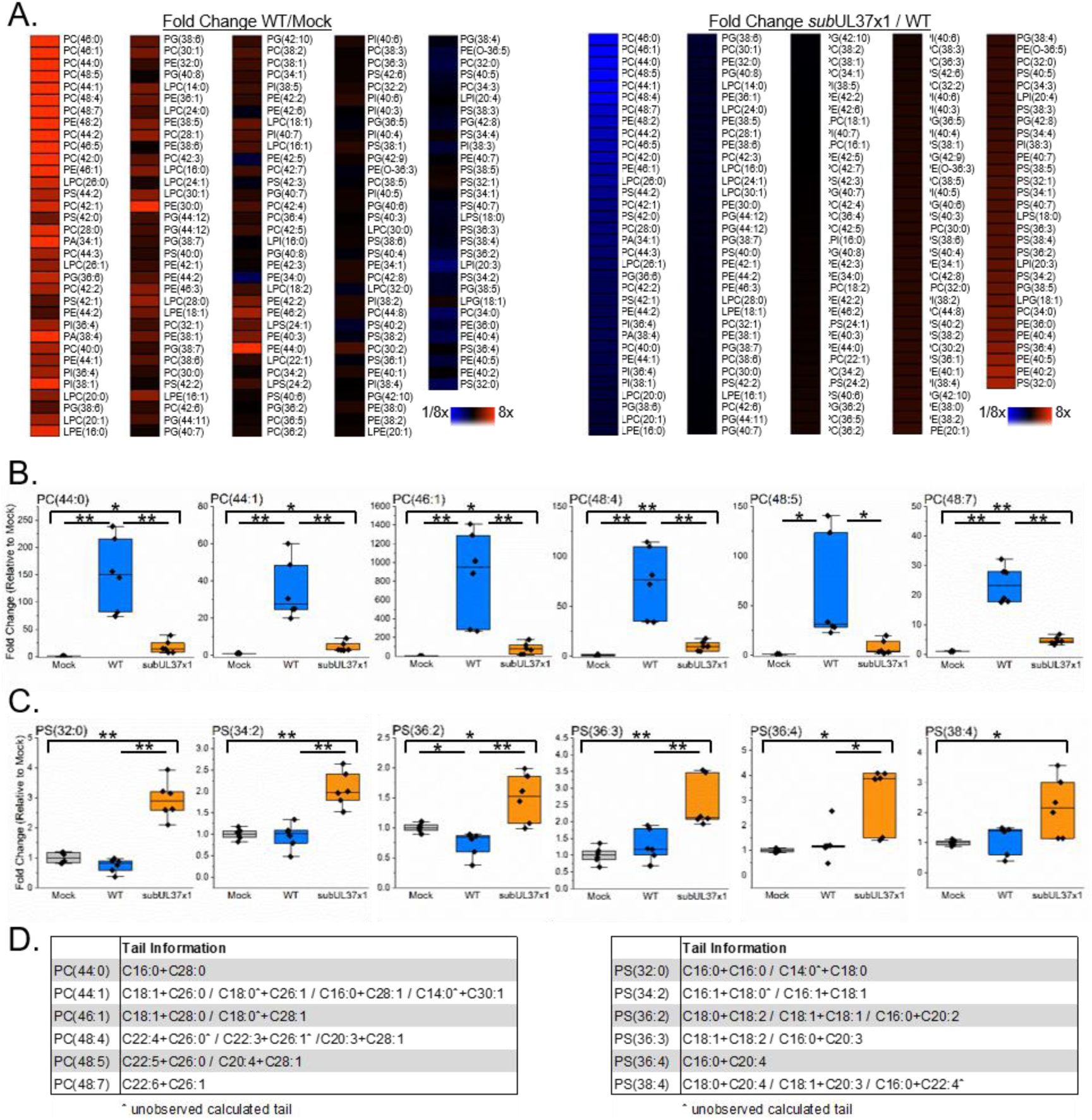
HCMV remodeling of host lipidome requires pUL37×1. (A, left) A LC-MS/MS phospholipidomic analysis of cells infected with subUL37×1 mutant virus compared to WT-infected cells. (A, right) The same lipidomic analysis comparing WT-infected to uninfected cells.(B) Phosphatidylcholine lipids (PC) with VLCFA tails altered by infection are shown. (C) Similar plots showing phosphatidylserine lipids (PS). (D) The FA tails in PC and PS lipids shown in (B-C) were identified by MS/MS. In some cases, only one of the two tails were observed. The non-observed tails are marked by a ^ symbol. All data is 3 dpi from cells infected at MOI = 3, N = >3.

When we examined phospholipids from *sub*UL37×1 infected cells, we observed a significant decrease in lipid levels compared to WT-infected cells (Fig. 9A, right panel). The most notably decreased lipids were the PC species that were greatly elevated by WT virus. All six of the PC species discussed above were significantly decreased in mutant-virus infected cells compared to WT (Fig. 9B). However, all species with the exception of PC(48:5) were elevated above the uninfected level by the mutant virus (Fig. 9B). PC lipids are important for membranes, including the virion envelope (31), suggesting that these lipids with VLCFA tails may be required for HCMV induced membrane reorganization and for virion infectivity.

Some lipids were increased by *sub*UL37×1 over WT levels (Fig. 9A). Several of these lipids were phosphatidylserine (PS) lipids. The mutant virus increased PS species by 4-fold or less (Fig. 9C), which was vastly less than the increases observed for PC lipids in WT infected cells (Fig. 9B). PS(36:2) was increased by only 1.5-fold. None contained a tail longer than C22 (Fig. 9D). Furthermore, none of these species were increased by WT virus and one—PS(36:2)—was slightly reduced in WT-infected cells compared to uninfected cells (Fig. 9C). These observations suggest that pUL37×1 suppresses their synthesis. Our lipidomics studies demonstrate that HCMV remodels the host lipidome—increasing the abundance of phospholipids with VLCFA tails—and that the magnitude of the lipidome shift requires pUL37×1.

HCMV activation of PERK is important for virus induced lipid metabolism (29). Both the level and activation of PERK are induced by HCMV (29, 49-51). Given our observation that pUL37×1 is important for HCMV lipidome remodeling and the release of Ca^2+^ from the ER may induce a stress response (52), we tested if pUL37×1 is required for HCMV-induced PERK protein levels. At 3 dpi, PERK levels were reduced 2-fold in mutant virus infected cells compared to WT-infected cells (Fig. 8). This observation suggests that pUL37×1 control of PERK may be important for HCMV lipid remodeling that we observed. Overall, our data demonstrate that pUL37×1 aids HCMV metabolic remodeling, particularly FA elongation and lipid metabolism.

## Discussion

Viral metabolic hijacking as a requirement for replication has been demonstrated in many diverse viruses (reviewed in (18, 19, 43, 44, 53-56)). The virus-host metabolic interaction is important to the infection of α-(13,15), γ-(57-59), and β-herpesviruses (11,12,20). Like HCMV, Kaposi’s sarcoma associated virus (KSHV) increases FA synthesis (60). Recently, the related γ-herpesvirus—murine gammaherpesvirus 68 (MHV68)—replication was suppressed by type I interferon induced limitations on fatty acid and cholesterol metabolism (61). These studies, along with those for HCMV and HSV (described in (18,43,44)) demonstrate that virus-metabolic interactions are essential to infection and can be targeted to limit replication.

HCMV globally remodeling of metabolism requires various host kinases (PERK (29), mTOR(12), CaMKK (22), and AMPK(21,32)) and transcription factors (SREBPs (12,28,33) and chREBP (62)). Our previous observation that HCMV induced FA metabolism requires mTOR activity lead to the finding that pUL38 is required for viral metabolic hijacking (12). We tested the hypothesis that a second viral protein pUL37×1 is required for HCMV metabolic remodeling. The results described here confirm our hypothesis that pUL37×1 also contributes to HCMV remodeling of host metabolism. Similar to pUL38, we found that pUL37×1 contributes to HCMV induced FA elongation and ELOVL7 expression (Fig. 4-6). Using our newly developed LC-MS/MS lipidomic methods, we found that pUL37×1 is needed for HCMV alteration of lipid metabolism (Fig. 9). During late stages of replication, pUL37×1 causes the accumulation of large vesicles (45). These vesicles may form due to the pUL37×1 dependent FA elongation and lipid synthesis described in this study. In addition to lipidome remodeling by HCMV, we found that pUL37×1 is also necessary for the virus to fully induce PERK expression (Fig. 8). This observation is consistent with the role of PERK in HCMV activation of lipogenesis (29). pUL37×1 may stimulate PERK expression by inducing an ER-stress response triggered when ER stored Ca^2+^ is released into the cytosol (52). Although at this time, we cannot rule out other possible mechanisms for pUL37×1 to contribute to the lipidome changes we observed.

Interestingly, our lipidomic studies further revealed that pUL37×1 suppresses PS lipid levels. PS lipids on the outer leaflet of the plasma membrane is an apoptotic signal (63,64). Our observations suggest that pUL37×1 is important in limiting the production of lipids associated with apoptosis. Since the PS species that are suppressed by pUL37×1 do not contain saturated VLCFAs produced by ELOVL7 (Fig. 9), it is likely that this function of pUL37×1 is independent of its ability to enhance FA elongation. PS synthesis occurs at the mitochondria-associated ER membrane in the same region where pUL37×1 is found (42) suggesting that it may act direction on PS synthesis enzymes. However, it is currently unknown if pUL37×1 interacts directly with the enzymes involved with PS synthesis. Importantly, our lipidomic observation suggest pUL37×1 further blocks apoptotic limitations on infection by reducing PS synthesis.

HCMV replication increases the flow of carbons from glucose to fatty acid and lipid synthesis (11, 24, 28,29, 33, 62). We have previously shown that glucose is metabolized to support fatty acid elongation (12). ELOVL7 is required for HCMV virus release and virion infectivity and synthesizes saturated VLCFAs that become part of the virion lipidome. Here, we report the robust increase of PL-VLCFAs by HCMV. Many of the PL-VLCFAs that were most dramatically increased by HCMV were PC lipids. PC species with shorter tails are the second most abundant PL in the HCMV virus envelope (31). Some of the PC-VLCFAs may be incorporated into the virion. Previously, we demonstrated that the loss of saturated VLCFAs results in a disruption of the infectivity per particle as measured by determining the ‘particle-to-infectious-unit ratio’ (12). Viruses produced by ELOVL7-depleted cells were 4-8-fold less infectious; i.e. a particle-to-IU ratio that of 400:1 to 800:1 compared to the WT ratio of 100:1 (12). *Sub*UL37×1 viruses also have a disruption in their particle-to-infectious-unit ratio (Fig. 2D). Our observations that *sub*UL37×1 virus has a disruption in FA elongation and lipid synthesis suggest that the loss of infectivity in the mutant virus is due—at least in part—to disruption in the virus-host metabolism interaction.

Our work establishes that pUL37×1 is an important player in virus-host metabolic interactions required for HCMV infection. This work demonstrates that HCMV employs multiple mechanisms—pUL37×1 as demonstrated here and pUL38 (12)—to ensure that the host metabolic activity is rewired. Furthermore, our results show that HCMV infection dramatically induces the accumulation of lipids with saturated VLCFAs and highlight the need to continue to study the role of lipid metabolism during HCMV infection. Based on our lipidomic studies, along with our metabolomic analyses (Fig. 3), we conclude that pUL37×1 is required for HCMV metabolic remodeling but is not necessary to induce viral metabolic hijacking. As such pUL37×1 and pUL38 likely coordinate the virus-host metabolism interaction along with other viral factors. Given the reliance of HCMV replication—and viruses in general—on metabolism it is important to define the viral and host mechanisms required for metabolic remodeling. Our results further support the critical functions of pUL37×1 in HCMV replication and provides a foundation for future studies in defining virus-host metabolism interactions to build upon.

## Materials and Methods

### Cells, Virus, and Experimental Setup

Human foreskin fibroblast cells (HFF) were cultured in DMEM containing 10% FBS, HEPES, and pen/strep. Prior to infection cells were maintained at full confluence for 3 days in serum-containing growth medium. Cells were switched to serum medium—DMEM, HEPES, and pen/strep—the day before infections or mock-infection. Cells were infected with either wild-type (WT) HCMV AD169 strain or *sub*UL37×1, a UL37×1-null virus (36). Both WT and mutant viruses were kindly provided by Dr. Thomas Shenk. All virus stocks were made by pelleting virions from the supernatant of infected cells through a 20% sorbitol. Viruses were resuspended in serum-free DMEM medium and stored at −80 °C. Infectious virus titers were measured by the visualizing cytopathic effect and the tissue culture infectious dose (TCID_50_) method. Since *sub*UL37×1 mutant virus shows minimal CPE, plaque identification was further confirmed by immunofluorescent microscopy. In this case, cells on the TCID_50_ plate were fixed using methanol and plaques were identified using an anti-pUL123 (IE1) monoclonal antibody (clone 1B12) followed by anti-mouse Alexa 488 secondary antibody. This method has similarly been used to quantify a mutant HCMV virus that lacks the UL38 gene (12). Particle-to-IU ratio was done using UL123-specific primers to measure viral DNA (65).

Metabolic tracer studies were done using 1,6-^13^C-glucose and fully labeled 1,2-^13^C-acetate sodium salt from Cambridge Isotopes (CLM-2717 and CLM-440, respectively). Following a 1 h infection period, cells were washed with warm PBS three times prior to the addition of serum-free DMEM medium containing either 4.5 g/L glucose (25 mM) or 0.0035 g/L acetate (60 µM). For 72 hpi samples, the labeling medium was replenished at 48 hpi.

### Protein Analysis

Proteins were examined by Western blot from whole-cell lysates prepared in reducing SDS-PAGE sample buffer (25 mM Tris pH 6.8, 5% SDS, 1% 2-mercaptoethanol, glycerol and complete protease inhibitor [Roche]). SDS-PAGE was performed in tris-glycine SDS running buffer using Mini-PROTEAN TGX gels (Bio-Rad). Proteins were transferred to Odyssey nitrocellulose membrane (Li-COR) using a wet-transfer system at 4 °C. Western blots were performed using 5% BSA in a Tris-buffered saline with 0.05% tween-20 (TBS-T) for blocking and antibody incubation, except for anti-ELOVL7, anti-ATP citrate lyase and anti-PERK blots which were performed using 3% milk in TBS-T. The following antibodies were used: mouse monoclonal anti-pUL37×1 clone 4B6-B (36), mouse monoclonal anti-pUL36 clone 10C8 (66), mouse monoclonal anti-pUL38 clone 8D6 (67), rabbit polyclonal anti-ELOVL7 (Sigma-Aldrich SAB3500390), rabbit monoclonal anti-AMPKα (Cell Signaling 2603), rabbit polyclonal anti-ATP citrate lyase (Thermo PA5-29497), rabbit polyclonal anti-ACC-1 (Cell Signaling 3662), rabbit polyclonal phospho-ACC-1 Ser79 (Cell Signaling 3661), rabbit monoclonal anti-PERK (Cell Signaling 3192), rabbit monoclonal anti-ACCS2 (Cell Signaling 3658), mouse monoclonal anti-actin (LI-COR 929-42212), and mouse monoclonal anti-α-tubulin (Sigma-Aldrich T6199). Blots using mouse monoclonal anti-HCMV, anti-actin and anti-tubulin antibodies were incubated for 1 h at room temperature. All others were done overnight at 4 °C. Quantification of Western blots was done using a LI-COR Odyssey CLx imaging system.

### Metabolomics and Metabolite Analysis

All metabolite measurements were done using liquid-chromatography high resolution tandem mass spectrometry (LC-MS/MS). At various times after infection, metabolic reactions were rapidly quenched using cold 80% methanol and extracted as previously described (14,23). Extracted water-soluble metabolites were dried under nitrogen gas and resuspended in Optima HPLC-grade water (Fisher Chemical) for reverse phase liquid chromatography or 50% methanol for hydrophilic interaction chromatography (HILIC). Mass spectrometric analysis was performed by four Thermo Scientific mass spectrometers: Quantum Ultra triple-quad, Quantum MAX triple-quad, Exactive orbitrap and Q-Exactive Plus orbitrap. Metabolites were measured in both positive and negative modes using reverse phase or HILIC separation (68,69). Metabolites were identified using mass spectral and UHPLC retention time data that were generated from external standards. Peaks were examined, and metabolites identified using Compound Discovery software (Thermo) and Metabolomic Analysis and Visualization Engine (MAVEN) (68,70). Each metabolite was measured using two technical replicates per experiment. Each metabolic experiment contained controls for determining ionization artifacts and sample degradation. All chemical and solutions for metabolomics, fatty acid analysis, and lipidomics. were HPCL or MS-grade.

### Fatty Acid Analysis

Fatty acid (FA) analyses were performed as previously described (12,17). Briefly, lipids were extracted from cells lysed using cold 50% methanol containing 0.05 M HCl. Cells were scraped and transferred to glass vials. Lipids were chloroform extracted and FA tails were chemical cleaved via saponification at 80 °C for 1h in a basic methanol buffer. The FAs were isolated using hexane and dried down under nitrogen gas. Following a resuspension in 1:1:0.3 methanol:chloroform:water or 1:1:1 methanol:chloroform:isopropanol FAs were analyzed by UHPLC and high-resolution mass spectrometry. FAs were examined using a Luna C8 reversed-phase column (Phenomemex) and Exactive orbitrap mass spectrometer (12) or a Q-Exactive Plus mass spectrometer. In the latter case, FAs were separated using the same column but utilized the same LC buffer solutions as used for lipid analysis described below. FA spectrums were collected using 140,000 resolution (at 200 m/z). Experiments utilizing metabolic tracers and FA labeling were additionally analyzed at 280,000 resolution. The data was analyzed and FA levels quantified using MAVEN. For labeling experiments, the data were corrected for naturally occurring carbon-13 using MATLAB (12). Each sample was measured using two technical replicates per experiment.

### Lipidomics and Lipid Analysis

Similar to the FA analysis, lipids for lipidomics were extracted by 50% methanol and chloroform. The chloroform was removed by drying the samples under nitrogen gas. Lipids were resuspended in 1:1:1 methanol:chloroform:isopropanol. If needed, samples were stored under nitrogen gas at −80 °C. Samples were normalized according to cell number: 125 µl of 1:1:1 resuspension solution per 100,000 cells. After resuspension, lipids were kept at 4 °C. Lipids were separated via UHPLC using a Kinetex 2.6 µm C18 column (Phenomenex 00F-4462-AN) at 60 °C on a Vanquish system. Two solvents were used for UHPLC: solvent A (40:60 water:methanol plus 10 mM ammonium formate and 0.1% formic acid) and solvent B (10:90 methanol:isopropanol plus 10 mM ammonium formate and 0.1% formic acid). UHPLC was performed at 0.25 ml/min flow rate using the following gradient conditions: 75% solvent A / 25% solvent B for 2 min; 35% A / 65% B for 2 min at curve value = 4; held at 35% A / 65% B for 1 min; 0% A / 100% B for 11 min at curve value = 4; and held at 0% A / 100% B for 4 mins. The column was briefly washed and re-equilibrated before the next run. In total each run lasted 30 min. All lipids were examined using a Q-Exactive Plus mass spectrometer operating in a data-dependent Full MS/dd-MS2 TopN mode. MS1 data was collected at either 70,000 or 140,000 resolution at 200 m/z using AGC = 1e6 with transient times of 250 ms and 520 ms, respectively. Spectrum were collected over a 200-1600 m/z mass range. MS2 spectrum were collected using 35,000 resoluton, AGC = 1e5 and 120 ms maximum injection time. Each sample was run twice, using 10 µl for negative mode and 8 µl of sample for positive mode. In negative mode, a normalized collision energy (NCE) value of 20 was used. In positive mode, the NCE value was increased to 30. All lipids were ionized using a heated electrospray ionization (ESI) source. The following ESI parameters were used in negative mode: sheath gas flow rate = 44; auxiliary gas flow rate = 14; sweep gas flow rate = 2; spray voltage = 3.3 kV; S-lens RF =76; aux gas temp 220 °C; and capillary temp = 320 °C. Similar settings were used in positive mode except the spray voltage was increased to 3.5 kV and S-lens RF level was decreased to 65. The instrument was calibrated weekly. For labeling experiments, the instrument was calibrated immediately before the start of analysis. As with metabolites and FA, lipids were measured using two technical replicates per experiment. Lipidomic data was analyzed using MAVEN and LipidSearch (Thermo Scientific).

## Acknowledgments

We acknowledge Thomas Shenk for providing guidance in our studies and reagents including the *sub*UL37×1 mutant virus and anti-HCMV monoclonal antibodies. We thank Joshua Rabinowitz for providing advice and assistance with the early stages of these studies. We are grateful to Felicia Goodrum, Jim Alwine, Lynn Enquist, the members of the Goodrum lab and the other members of the Purdy lab for helpful discussion regarding this work.

This work was funded in part by a grant from the Arizona Biomedical Research Commission as made available through the Arizona Department of Health Services to J.G.P. Additional funding was provided by the BIO5 Institute, the Department of Immunobiology, the College of Medicine-Tucson at the University of Arizona, and a New Investigator Award from the University of Arizona Health Science award to J.G.P.

